# MIA: An Open Source Standalone Deep Learning Application for Microscopic Image Analysis

**DOI:** 10.1101/2022.01.14.476308

**Authors:** Nils Körber

## Abstract

In recent years the amount of data generated by imaging techniques has grown rapidly along with increasing computational power and the development of deep learning algorithms. To address the need for powerful automated image analysis tools for a broad range of applications in the biomedical sciences, we present the Microscopic Image Analyzer (MIA). MIA combines a graphical user interface that obviates the need for programming skills with state-of-the-art deep learning algorithms for segmentation, object detection, and classification. It runs as a standalone, platform-independent application and is compatible with commonly used open source software packages. The software provides a unified interface for easy image labeling, model training and inference. Furthermore the software was evaluated in a public competition and performed among the top three for all tested data sets. The source code is available on https://github.com/MIAnalyzer/MIA.

## 1 Introduction

Convolutional deep neural networks are state-of-the-art for most computer vision tasks and often outperform classical image analysis techniques [1]. Neural networks are trained by error backpropagation, i.e. they learn based on pairs of an image input and a target network output (ground truth), minimizing the error between the network output given the input and the ground truth [2]. The features of the network that map input to output are learned during training and ideally generalize to correctly identify unseen objects of a learned object class. Consequently, deep learning requires minimal domain knowledge as no feature engineering is required making the same network architecture applicable for the identification of completely different objects and imaging modalities. In practice this means, instead of creating a sequence of operations (e.g. background subtraction, filtering, thresholding, separation by object size) as used in classical image processing, the only inputs to a deep neural network are an image and the corresponding target objects to be detected. The underlying features that identify the target objects within an image are learned from the training data. Over the last years, networks with growing depth [3], increasing complexity [4, 5, 6] or expanding number of parameters [7] have been designed for computer vision tasks such as classification [8], semantic segmentation [9] or object detection [10]. While the performance of a network usually increases with the amount of training data, so-called transfer learning reuses the model weights of a network trained with a large general dataset to achieve generalization on the target dataset with less data [11].

In recent years, deep learning has slowly evolved from the field of computer science into applications in microscopy, achieving impressive results in areas such as cancer detection [12, 13], super-resolution [14], denoising [15, 16] or label-free staining [17, 18]. With the massive growth of image data generated by microscopy and high-throughput imaging, the demand for automated image analysis solutions has increased drastically. So far, the implementation and training of deep learning algorithms for image analysis required advanced coding skills, mathematical expertise as well as the computational resources, which limited its broad application in the life science community. Several solutions have been deployed to reduce these hurdles and provide an easily accessible interface for deep learning, such as DeepCell Kiosk [19], ImJoy [20], ZeroCostDL4Mic [21], U-Net plugin [22], ilastik [23], CellProfiler [24], 3DeeCellTracker [25], and CDeep3M [26]. While each of these applications provide great value for certain aspects of the deep learning procedure, like training, cloud computation, or pre-trained model integration, a comprehensive, unified, and user-friendly tool that integrates these features into a single versatile software is lacking.

Here, we developed MIA - the Microscopic Image Analyzer - as a standalone software with a graphical interface integrating image labeling, model training and inference that can be operated platform-independent (figure 1). MIA was designed with a particular focus on usability and simplicity. Although the model training time is often considered as critical, the most laborious part of the deep learning procedure is by far image labeling. Assisted labeling can significantly reduce the required hands-on working hours. Whereas comparable applications require expert knowledge or have very limited access to complex features, MIA offers them as selectable built-in options. Users can choose from a number of neural network architectures, with several training options providing experienced users with the option to customize for a specific data set while still achieving good results with default settings. The training computation can be performed on the graphical processing unit (GPU) or the central processing unit (CPU) depending on the available hardware. The training process and model performance are monitored during training and the prediction of unseen data can be visually inspected and corrected if necessary. Finally, the results can be saved for individually detected objects and labels can be exported for further analysis with other software.

**Figure 1:**
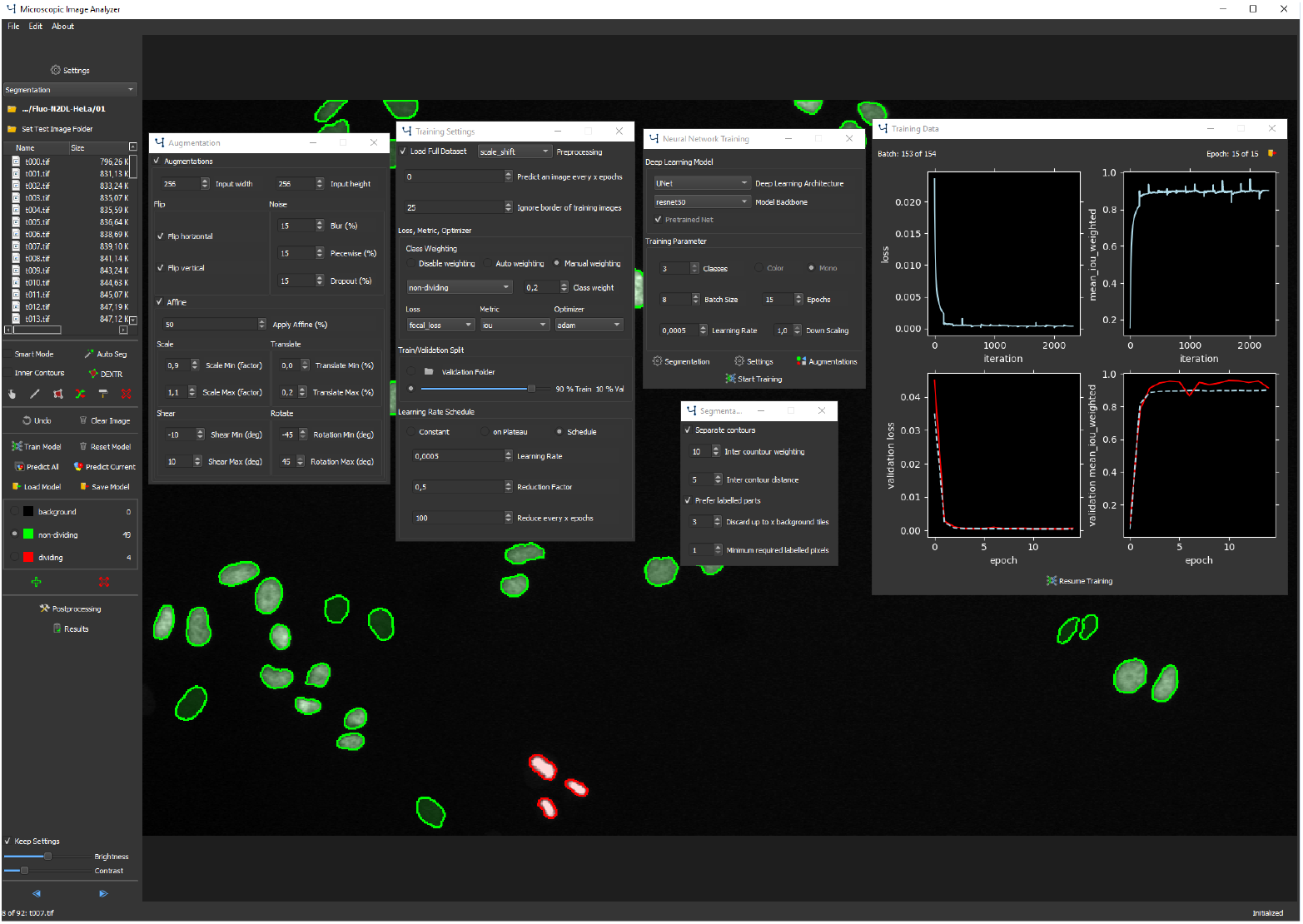
The MIA user interface. The left panel of the user interface shows an explorer interface for file navigation, tools for image labeling and options for model training, prediction, saving, loading, post-processing as well as setup of object classes and other options. The center panel shows the currently displayed image with labeled cells, overlaid with several open windows showing training options, augmentation settings, training progress and others. The currently displayed image shows HeLa cells expressing H2b-GFP [27].

## 2 MIA Features

The motivation for the development of MIA was to create a tool that gives any user access to powerful deep learning image analysis in order to address different types of scientific questions. However, to cover a wide range of applications and imaging modalities, training parameters and technological depth still need to be adjustable to achieve excellent results for each use case. Since easy access to deep learning competes with freedom and performance, MIA lowers the entry-level for deep learning at the expense of flexibility. See table 1 for a comparison of MIA with other methods used in microscopic image analysis.

**Table 1:** Comparison of MIA with classical image analysis and deep learning

| Requirements | Classical Analysis | Deep learning | MIA |
| --- | --- | --- | --- |
| Apriori domain knowledge | medium–high | low | low |
| Training data | none–few | medium–high | medium–high |
| Training/processing time | low | high | high |
| Coding skills/mathematical expertise | low–medium | high | none |
| Development effort | low–medium | high | low |
| Image labeling tools | not required | external | included |
| Capabilities | Classical Analysis | Deep learning | MIA |
| Improves with increasing data | no | yes | yes |
| Generalization | low | high | high |
| Performance | low–medium | high | high |
| Flexibility | medium | high | medium |
| Application range | medium | high | medium |
| Revision of predictions | external | external | included |

The scope of MIA is the analysis of microscopic data. Therefore image classification, semantic segmentation and object detection have been implemented along with a tracking mode as post-processing of previously detected objects to cover the most common tasks. To test and demonstrate the features of MIA, training datasets for classification, segmentation, and object detection were generated based on the same image source showing HeLa cells expressing H2b-GFP [27] that were categorized in dividing and non-dividing cells. In the following sections these datasets were utilized to showcase different applications of MIA.

### 2.1 Image Classification

Image classification is the categorization of each image to a predefined class. For example, it can be used to identify different cell types or different phenotypes following drug treatment. Since a complete image is assigned to only one single class label, the information where in the image a particular object is located is not considered, yet the labeling process is much faster compared to labeling of individual pixels. For the classification with MIA, the user can choose from a variety of implemented network architectures ranging from fast networks like mobilenet [28] to larger and slower models like NASNet [29]. See https://github.com/MIAnalyzer/MIAfor all available model architectures and additional details.

To showcase the capabilities of MIA for classification, a neural network was trained with default settings on a dataset with dividing and non-dividing HeLa cells expressing H2b-GFP [27]. Each image of the training set consists of a small image patch containing the target cell and a corresponding label that categorizes the cell into dividing or non-dividing (figure 2A). The trained model achieved more than 95 % accuracy on unseen images to correctly classify them as dividing or non-dividing. Note that the model was able to identify cells in prophase of mitosis, some of which are barely distinguishable from cells in interphase.

**Figure 2:**
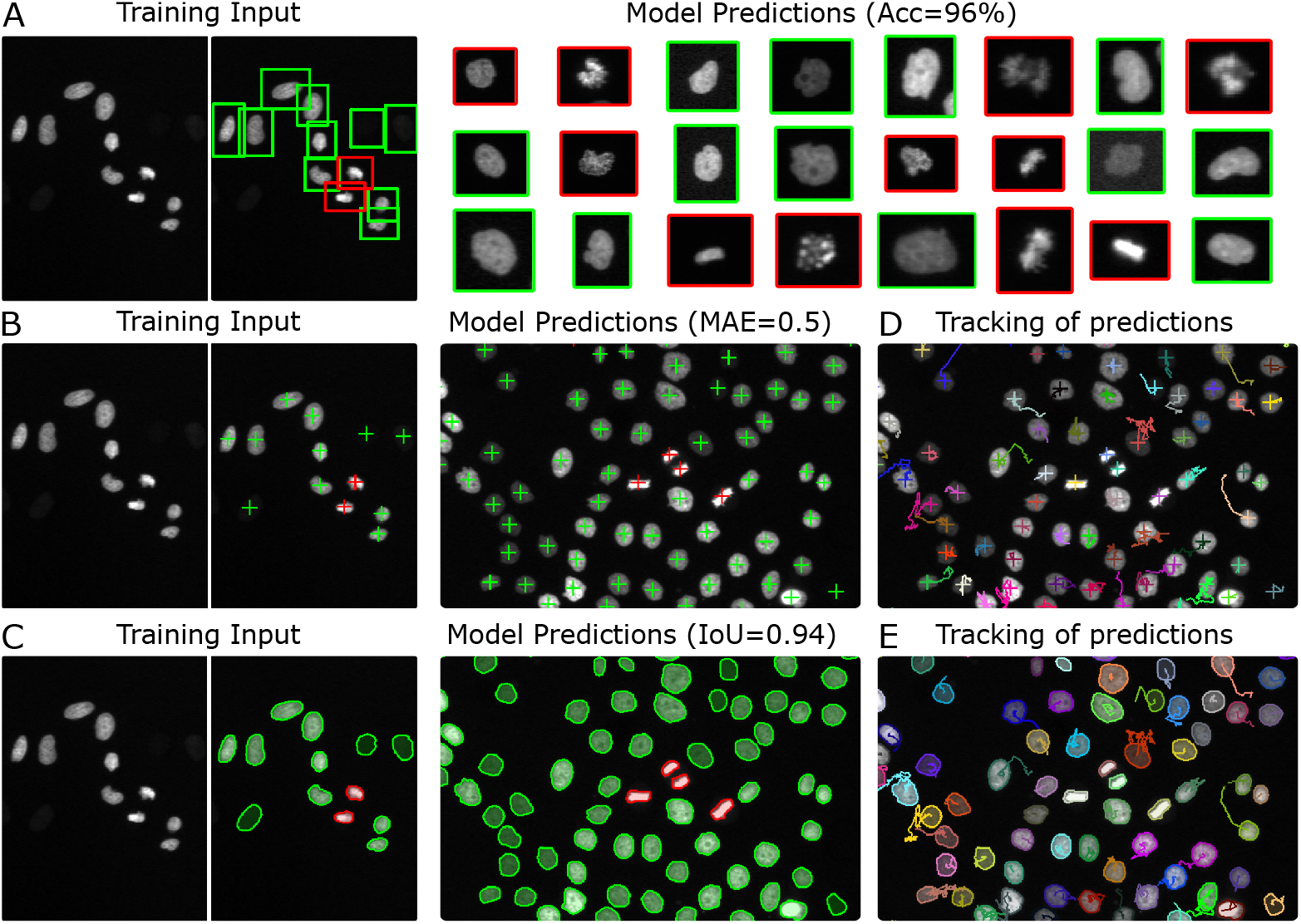
Example applications using MIA. Neural networks were trained for classification (A), object detection (B), and semantic segmentation (C) using the same dataset of HeLa cells expressing H2b-GFP [27]. Cells were categorized in dividing (red) and non-dividing (green) cells. Training input showing a training image (left) along with overlaid corresponding labels (right), and model predictions showing the model output as an overlay of the source image with predicted labels are displayed. A) A neural network was trained for classification of dividing and non dividing cells. The training images were split into patches that contain a single target cell as neural network input. The model achieved an accuracy (Acc) of 96 % on the validation set to correctly classify an image as dividing or non-dividing. B) A neural network was trained for object detection using the same training set, preserving the position information of target objects, achieving a mean absolute error (MAE) of 0.5. C) A neural network trained for semantic semantic segmentation was trained to predict pixelwise class labels with an intersection over union (IoU) of 0.94 on the validation set. D, E) Results from object detection (D) or segmentation (E) were used to perform tracking of individual objects over consecutive time points. Each object is shown in an individual color and traces refer to positions along previous time points.

### 2.2 Semantic Segmentation

Semantic segmentation is the process of assigning a predefined class to every pixel of an image. It allows foreground and background as well as different cell or tissue types to be distinguished. The number of classes need to be defined prior to training, and all objects need to be labeled with their corresponding class with the unlabeled area assigned to background. MIA provides several labeling tools for semantic segmentation, such as polygon tool, freeform tool, slicing tool, an extension, and an erase tool. However, to label objects one by one can be tedious for complex structures and large amount of objects. Hence, automated or semi-automated labeling tools have been implemented based on Grabcut [30], Holistically-nested edge detection [31], and Deep extreme cut [32], that may reduce labeling time drastically [32]. Several neural network architectures for semantic segmentation including the popular U-Net [33] or DeepLab [34] can be chosen building on a variety of selectable model backbones from computational less expensive models like mobilenetv2 [28] to more complex models like EfficientNet [35]. See https://github.com/MIAnalyzer/MIAfor all available model architectures and additional details. To avoid fusion of neighboring objects as a common problem for segmentation of dense microscopic images, MIA supports increasing the weighting of pixels in close proximity to adjacent objects, as shown in [33]; in that way, the network learns to separate neighboring objects during training.

To illustrate the applicability of MIA for semantic segmentation, a U-Net [33] was trained using default settings to segment HeLa cells expressing H2b-GFP [27]. The training images consist of cells and corresponding pixel-precise labels outlining dividing and non-dividing cells (figure 2C). The trained model achieved a mean intersection over union of 0.94 for unseen data to correctly classify pixels to the corresponding class label. Shape, size and position of each dividing and non-dividing cell can be extracted from the predicted image data.

### 2.3 Object Detection

Object detection identifies all instances of all classes and their respective position in the image. Its primary task is to count objects, for example, cells of different cell types in a tissue and their spatial distribution. In computer vision, object detection is often defined as the detection of each object instance along with its corresponding bounding box. However, in MIA, we designed object detection by the center position of the object omitting the bounding box, as in the life sciences the bounding box rarely offers relevant information. To measure the size of detected objects, it is recommended to use semantic segmentation instead. For object detection in MIA, the same model architectures can be chosen as for semantic segmentation, but a linear layer is used in the final layer of the network instead of a softmax or sigmoid layer, enabling the regression of peak values at image spots where objects are located.

To exemplify the utilization of object detection, MIA was used with default settings to train a model for the detection of dividing and non-dividing HeLa cells expressing H2b-GFP [27]. The dataset comprises of training images of cells and corresponding labels with the position information of dividing and non-dividing cells (figure 2B). The trained model achieved a mean absolute error of 0.5 on unseen images to correctly identify localizations of target cells. Exported results can be used for further analysis, e.g. to describe cell division events as a function of cell density.

### 2.4 Object Tracking

Object tracking is the identification of the same object at consecutive time steps. One application of object tracking is to determine the trajectories of living cells in tissue or during development. Tracking is implemented as a post-processing step and can be used in combination with object detection or semantic segmentation. The center of each object is calculated and compared to the centers in the subsequent frame by minimizing the total distance between the objects in the two frames based on the Hungarian algorithm [36]. An object that is not detected in several consecutive frames is considered lost. In addition to automatic tracking, MIA allows to reassign detected instances to different objects to correct for potential errors. Furthermore, MIA supports tracking of different object types in parallel.

To illustrate object tracking with MIA, the results for dividing and non-dividing HeLa cells from semantic segmentation and object detection were used to calculate the position of each object in a time sequence (figure 2D,E). Based on the detected objects in the image sequence, the detections of the same object in sequential images were combined to obtain the position information of that object along the entire sequence. As a result, motion profiles of unique objects with respect to the image sequence could be identified. The results may be used for further analysis, to calculate, for example, the cell movement in relation to division events.

## 3 Recommended MIA Workflow

MIA supports input data from all common public image types and formats including 16-bit depth images and z-stacks. The images can be labeled according to the respective target application, i.e. classification, semantic segmentation or object detection. The initial decision which objects to label and which classes of objects to choose have a significant impact on the final result and depend on the specific scientific question. Once images are labeled, the training process can be started by selecting a model architecture. All models can be trained from scratch or, alternatively, using pre-trained weights, which were derived from training on a large dataset like imagenet [38]. This so-called transfer learning allows faster convergence for smaller datasets. Another way to increase the amount of training data is data augmentation, i.e. modifying the images of the training set by image operations like flipping, translation or scaling. MIA provides a number of possible augmentations that are automatically applied to the input images during training.

Hyperparameters, such as batch size or a specific learning rate schedule, can be defined via the user interface. For model optimization, the most commonly used loss functions, e.g. cross-entropy, and optimizers, e.g. Adam [39], can be selected depending on the particular application. To monitor the model performance, it is common practice to split the training set into a training and a validation set, which can be done in MIA either randomly or by specification of a data subset. Real world datasets are usually not balanced well, meaning that the frequency of different objects can differ strongly within the dataset. To circumvent this class imbalance, several options are implemented in MIA such as automatic class weighting, removal of less labeled image parts during training or specific loss functions like focal loss [40].

Once the model training is finished, the trained model can be applied to predict unseen images. The inference is always specified by the selected application, e.g. a model trained for semantic segmentation yields a model that can be used to segment images that are similar to the images present in the training set. All predicted images can be visually inspected and corrected if objects were incorrectly classified.

We propose the following workflow to create a neural network performing well on a real world dataset (figure 3):

**Figure 3:**
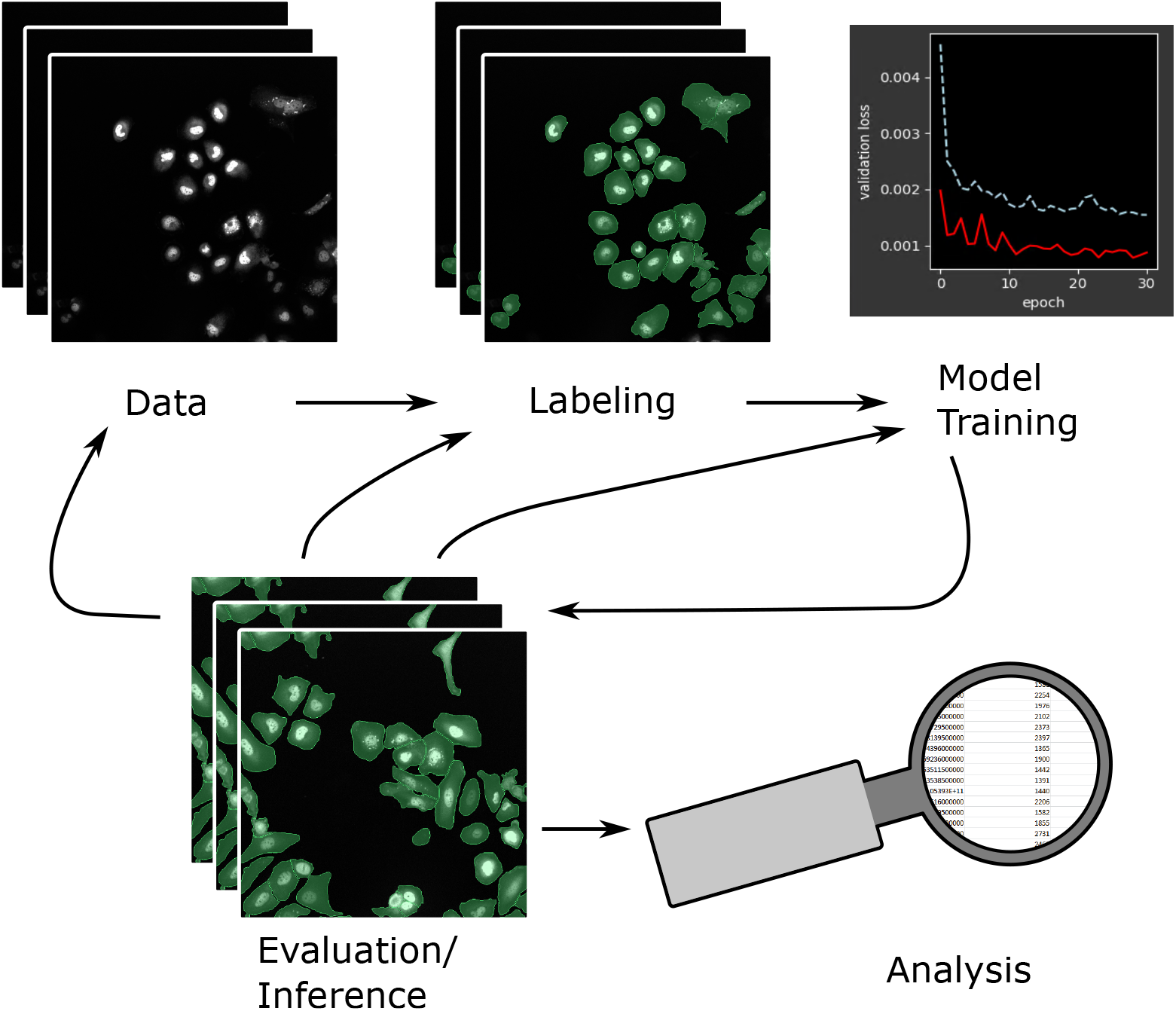
Proposed workflow for efficient neural network training. Start with image labeling, train a model and evaluate its performance. From there either label new data, correct predicted labels or train a new model. Repeat this process until a sufficiently performing model is trained. Finally, the model can be used for prediction and analysis. Images show human hepatocarcinoma-derived cells expressing YFP-TIA-1 [37].

1. Label a small set of images.
2. Train a model.
3. Repeat until the model performs sufficiently well on unseen data:
  a. Predict unlabeled images and evaluate model performance.
  b. 
    i. Correct the predicted image labels,
    ii. label additional images,
    iii. or skip this step.
  c. Update training dataset and retrain the model.

**Note:** training can be continued with conserved weights from previous training or started from scratch.

A significant advantage of this iterative approach with a human-in-the-loop is that only rare cases need to be corrected, whereas correctly predicted labels can be reused without further work. This highly reduces the redundancy of the data and a large diversity of the dataset is granted. Thus, the labeling is focused on data that need supervision, whereas other parts for which the model is already performing well can be ignored.

All trained models can be saved, reloaded and further trained. Stored models always create identical predictions for the same input, so they can be used for archiving and quality control. Finally, detected objects can be analyzed based on their size, shape, position or signal intensity, and they can be exported for further analysis.

## 4 Cell Segmentation Benchmark

In order to evaluate the performance of MIA, the software was tested on datasets being part of the Cell Tracking Challenge, a public competition for image analysis algorithms [41]. The performance of MIA was compared to the state-of-the-art results of the Cell Segmentation Benchmark (CSB). The challenge offers different datasets derived from 2D or 3D time series experiments or simulations. Each dataset consists of a training set and a test set. The training set contains images and corresponding labels for segmentation, whereby the correctness and completeness of the labels varies for each dataset. The test set only comprises images while the corresponding labels remain with the challenge organizers and are used for evaluation. The evaluation is based on the mean of an object detection score and an object-based intersection over union; see challenge website for details [41]. MIA was evaluated using three datasets each showing a different use case and relation to a real world application.

The first two datasets used for analysis with MIA contained complete but partly incorrect label information. The first dataset (FLUO-N2DH-SIM+ [42]) consists of simulated HL60 cells stained with Hoechst. Due to the nature of a simulated dataset, the image labels are exact and complete for the images of the training set. The second dataset (Fluo-N2DL-HeLa [27]) consists of images of HeLa cells stably expressing H2b-GFP. The corresponding labels are computer generated from previous challenge submissions. In that way, all training images have a corresponding label, but labels may be incorrect for difficult cases. In order to enhance the quality of these labels, MIA was used to quickly screen through the data and all labels which were obviously incorrect, i.e. missing a part of a target cell, were corrected by hand. For these two datasets, a DeepLabv3+ [34] model architecture with a Xception backbone [43] was chosen and trained on the training set. The trained models were used to evaluate their performance on the test set and achieved 2nd and 3rd rank of all submissions without bells and whistles (table 2).

**Table 2:**
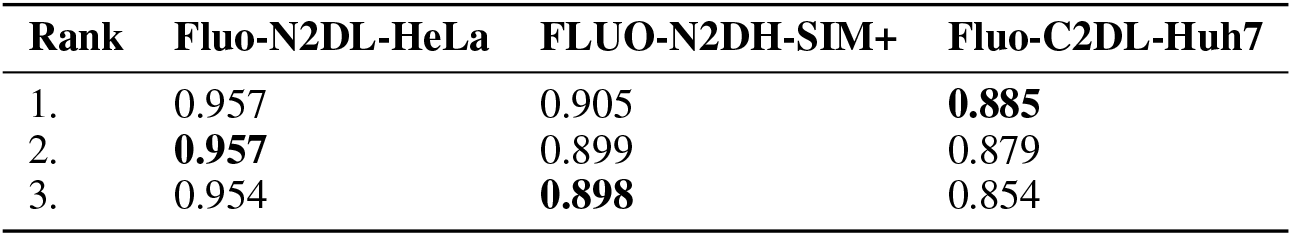
Top three results of the Cell Segmentation Benchmark (OP_CSB_) [41] for 3 datasets. The top three results of the CSB are shown with MIA results in bold, see http://celltrackingchallenge.net/participants/BFR-GE/.

| Rank | Fluo-N2DL-HeLa | FLUO-N2DH-SIM+ | Fluo-C2DL-Huh7 |
| --- | --- | --- | --- |
| 1. | 0.957 | 0.905 | **0.885** |
| 2. | **0.957** | 0.899 | 0.879 |
| 3. | 0.954 | **0.898** | 0.854 |

However, real world images usually have no inherently labeled objects. To further evaluate MIA in a more realistic scenario, the Fluo-C2DL-Huh7 dataset of human hepatocarcinoma-derived cells expressing the fusion protein YFP-TIA-1 [37] was used. This training data consists of 58 images, from which only 13 are labeled. Furthermore, the difficulty of correctly segmenting the cells is higher compared to the other datasets because the cells are very dense and their boundaries are visually indistinct. To tackle this task, the workflow from the previous section was applied. A small model was trained on the existing 13 labeled images intentionally overfitting to the training data. This model was then used to predict the labels for the remaining unlabeled images of the training set. These predicted labels were corrected by hand utilizing MIA to create a complete training set. In addition, the task was treated as a three-class problem defining background, target cells and the outer border of cells (three pixels wide) as individual object classes. Adjacent cells can be separated more easily when borders of cells are treated as an additional class since the network needs to explicitly identify the border region of cells. Finally, a U-Net [33] with an Inception-v4 [44] backbone was trained on the training data. With this approach, MIA achieved the new state-of-the-art for this dataset (table 2).

## 5 Discussion

With the development of MIA, we aim to simplify the use of deep learning for image analysis, with a particular focus on the diverse field of microscopy. MIA enables users without any programming experience to perform state-of-the-art image analysis based on deep learning for a wide range of applications providing all necessary tools in one single application. Furthermore, the access to hyperparameters and training options can improve the understanding of deep learning methods and strengthen their acceptance. As MIA runs on local hardware, it is independent of any cloud service and can be scaled based on resources and requirements.

We found that most existing applications focus on a specific part of the deep learning sequence, such as training, inference or a specific solitary task. On the other hand, a single application comprising all processing steps minimizes errors in data transfer and facilitates a seamless and optimized data flow. With MIA, we attempted to integrate the complete process, proposing a workflow with the human-in-the-loop that is fast and flexible to create powerful models for most real world applications. Default parameters in MIA depict the most commonly used options, thus inexperienced users are able to successfully train a model on any dataset, while users experienced in deep learning have all the options to customize the analysis to achieve state-of-the-art results.

One major focus of MIA is image labeling, which is often not integrated in deep learning solutions, but at the same time often is the most time-consuming part for the user in practice. MIA supplies automated solutions for image labeling that reduce hands-on time and the frustration of repetitive work.

With the current version of MIA, it is possible to perform classification, object detection, segmentation as well as tracking, which covers a broad range of the microscopy image analysis portfolio. Future development could include the implementation of instance segmentation, 3D models or automated hyperparameter tuning.

All predictions generated by MIA can be exported as image mask for subsequent use with other image analysis software such as ImageJ [45] or CellProfiler [24], placing the focus on deep learning while leaving downstream analysis or a larger image processing pipeline to external software. Along with this, trained models can be saved and reused with other applications or custom code using the tensorflow SavedModel format [46], making the models portable and compatible for model exchange in accordance with FAIR data principles [47].

In summary, we provide a versatile tool that allows an easy access to state-of-the-art deep learning based image analysis for a broad range of applications.

## 6 Methods

### 6.1 Implementation Details of MIA

#### 6.1.1 Environment

MIA is entirely written in Python using several publicly available open source solutions. The graphical user interface is written with PyQt (Riverbank Computing), the image processing and model computation are based mainly on Numpy [48], OpenCV [49], scikit-image [50], imgaug [51], TensorFlow [46] and Keras [52]. The model architectures are relying on open source repositories [53, 54, 55, 56] and custom implementations. The complete source code of MIA can be found on https://github.com/MIAnalyzer/MIAwith a user manual, installation options and additional information.

#### 6.1.2 Requirements

MIA is cross-platform compatible and has been tested successfully on several systems running with different versions of Windows and Linux on workstations or servers. The software supports GPU acceleration of most CUDA [57]-compatible units. Even though training can be performed with a CPU, it is highly recommended to use a GPU to decrease training time significantly.

### 6.2 Datasets for the Demonstration of MIA Features

To showcase applications using MIA, different datasets for neural network training were generated derived from the same images showing HeLa cells expressing H2b-GFP [27]. The silver segmentation labels provided by the Cell Tracking Challenge [41] were used as starting point for label generation. MIA was used to introduce an additional third class to this binary labeled data. All dividing cells, i.e. cells that were split into two objects in the subsequent time point and two objects deriving from a single object from the previous time point, were reassigned to an extra class label. All remaining objects were kept unchanged, resulting in segmentation labels for dividing and non-dividing cells.

Labels for object detection were generated from these segmentation labels by calculating the center of each contour using image moments.

Training pairs for image classification were generated by creating image patches and their associated labels based on the bounding box of each segmented dividing and non-dividing cell extended by 12 pixels in all directions, whereby only every tenth image of a non-dividing cell was used.

Training of the neural networks for these datasets was performed using MIA’s default settings for the target application. The validation data for each application was created by randomly withholding 15 % of the training data from the training.

Note that the evaluation metrics, shown in section 2, are based on the validation data and may differ from potential values of the test data, since the test data are from different experiments and the corresponding labels are unknown. Predictions of images shown in figure 2 belong to the test data.

### 6.3 Participation in the Cell Segmentation Benchmark

All steps used to participate in the Cell Tracking Challenge were done with the source code of MIA, except the conversion of the Cell Tracking Challenge labels to MIA compatible labels and vice versa. Since MIA counts touching objects as one, unlike the challenge, closely connected objects were separated by a zero-valued boundary between them when converted to MIA and expanded to the adjacent objects when converted back to the challenge labels to create touching objects.

All models were trained with 512×512 image patches that are randomly sampled from the training images. The same augmentation strategy was used for all trainings using image flipping, rotation, up to 10 % shearing, 90–110 % scaling and a 15 % probability of image blurring, piecewise affine image transformation or image dropout. The Adam optimizer [39] with an initial learning rate of 0.0005, halved every 50 epochs, was used to optimize the cross entropy cost function.

Fluo-N2DH-SIM+ [42]: A DeepLabv3+ [34] with Xception [43] backbone model was trained from scratch with a batch size of 16 for 500 epochs. The target cell class was weighted with 0.6 and background with 0.4 accordingly. Pixels between objects were weighted with a maximum weighting of *ω*_0_ = 10 and a border distance parameter of *σ* = 5 (as calculated in [33]).

Fluo-N2DL-HeLa [27]: A DeepLabv3+ [34] with Xception [43] backbone model with pre-trained weights was trained with a batch size of 16 for 300 epochs, using a class weighting of 0.9 to 0.1, a maximum inter-object pixel weighting of *ω*_0_ = 28 with a border distance parameter of *σ* = 5 (as calculated in [33]).

Fluo-C2DL-Huh7 [37]: A U-Net [33] with an Inception-v4 [44] backbone with pre-trained weights was trained with a batch size of 8 for 500 epochs. The cell border class was weighted with 1, the center cell class with 0.1 and background with 0.05.

All model predictions were done with test time augmentation, meaning that each input image was flipped and rotated by ±90 degree to generate a total of six input images. The model predictions of these transformed images were averaged to the final prediction. Post-processing to separate erroneously connected objects was either omitted or the detected objects were split based on watershed algorithm [58] using the predicted class probabilities or based on morphological image operations.

## 7 Acknowledgment

I would like thank my colleagues at BfR for their help and scientific input, in particular Chris Tina Höfer, Sebastian Dunst, Mariana Lara Neves, and Gilbert Schönfelder for critical comments on the manuscript. Furthermore, I thank Tra My Tran as the first user of the software for providing feedback and reporting bugs.

